# Stretching and squeezing of neuronal log firing rate distribution by psychedelic and intrinsic brain state transitions

**DOI:** 10.1101/2021.08.22.457198

**Authors:** Bradley Dearnley, Martynas Dervinis, Melissa Shaw, Michael Okun

## Abstract

How psychedelic drugs change the activity of cortical neuronal populations and whether such changes are specific to transition into the psychedelic brain state or shared with other brain state transitions is not well understood. Here, we used Neuropixels probes to record from large populations of neurons in prefrontal cortex of mice given the psychedelic drug TCB-2. Drug ingestion significantly stretched the distribution of log firing rates of the population of recorded neurons. This phenomenon was previously observed across transitions between sleep and wakefulness, which suggested that stretching of the log-rate distribution can be triggered by different kinds of brain state transitions and prompted us to examine it in more detail. We found that modulation of the width of the log-rate distribution of a neuronal population occurred in multiple areas of the cortex and in the hippocampus even in awake drug-free mice, driven by intrinsic fluctuations in their arousal level. Arousal, however, did not explain the stretching of the log-rate distribution by TCB-2. In both psychedelic and naturally occurring brain state transitions, the stretching or squeezing of the log-rate distribution of an entire neuronal population reflected concomitant changes in two subpopulations, with one subpopulation undergoing a downregulation and often also stretching of its neurons’ log-rate distribution, while the other subpopulation undergoes upregulation and often also a squeeze of its log-rate distribution. In both subpopulations, the stretching and squeezing were a signature of a greater relative impact of the brain state transition on the rates of the slow-firing neurons. These findings reveal a generic pattern of reorganisation of neuronal firing rates by different kinds of brain state transitions.

## Introduction

Elucidating the physiological processes through which psychedelics impact brain activity is motivated both by the need to advance our basic understanding of the different states of wakefulness and consciousness (Vollenweider and Geyer, 2001; Foster et al., 2016; Brecier et al., 2021) and by the therapeutic potential of this class of compounds (Carhart-Harris et al., 2016b, 2017; Johnson and Griffiths, 2017; Roseman et al., 2018; Carhart-Harris et al., 2021). At the cellular level, psychedelics exert their effect primarily by activating specific intracellular signalling pathways via the excitatory G-protein coupled serotonin-2A receptors (5-HT_2A_Rs, e.g., see Nichols, 2016, Madsen et al., 2019). Recent fMRI, MEG and EEG studies in human subjects uncovered profound alterations in macroscale brain resting state activity and functional connectivity after psychedelics administration. These alterations included increased functional connectivity between nodes of the default mode network and reduced brain-wide low frequency power (Carhart-Harris et al., 2012, 2016a; Muthukumaraswamy et al., 2013; Preller et al., 2018, 2019; Timmermann et al., 2019; Luppi et al., 2021), and were suggested to be directly driven by a change in the excitability of individual neurons, caused by activation of their 5-HT2ARs (Deco et al., 2018; Burt et al., 2021). However, this suggestion remains largely untested, as the effects of psychedelic drugs on the cortical microcircuit activity have received only limited attention and remain largely unknown. Indeed, only a few studies investigated the changes in spontaneous and sensory evoked activity in cortical neurons after systemic administration of a psychedelic drug in non-anaesthetised mammals (Wood et al., 2012; Gener et al., 2019; Michaiel et al., 2019), and they were limited to recording multi-unit activity or a relatively low number of neurons per animal. Here, we utilised the ability to simultaneously record from hundreds of individual neurons, afforded by the Neuropixels probes (Jun et al., 2017), to study how an orally delivered psychedelic compound changes the activity of neuronal populations in mouse medial prefrontal cortex (mPFC). We focused on mPFC since its neurons have a high level of expression of 5-HT2AR (Santana et al., 2004; Weber and Andrade, 2010; Andrade, 2011), it is implicated in the functional connectivity changes described in human subjects and it has been a primary focus of previous in vitro research (Avesar and Gulledge, 2012; Halberstadt, 2015).

Consistent with previous studies, we observed that a psychedelic drug produced a bidirectional modulation of firing rates, with rates of large neuronal subpopulations being elevated and suppressed. Furthermore, by relying on the high yield of single units in our recordings, we made the novel observation that this bidirectional modulation results in a prominent and robust increase in the width of the log firing rate distribution of the neuronal population. Previously, modulation of the width of the log-rate distribution was observed across sleep-wakefulness transitions (Watson et al., 2016). Since transition into a psychedelic brain state is distinct from changes along the sleep-wakefulness axis, our observation suggested that modulation of the width of the log-rate distribution is not unique to transitions between different sleep states, and prompted us to consider its relationship with brain state in more detail. We found that changes in pupil-indexed arousal in awake, drug-free mice are also correlated with changes in the width of log-rates in mPFC and other brain regions. We show that both psychedelic and arousal driven changes are in fact instances of a general phenomenon of stretching and squeezing of the log-rate distribution of an entire neuronal population as a result of the existence of two prominent subpopulations of neurons whose firing rates are modulated in opposite directions. Often the log-rate distribution of a subpopulation that is shifted to the right (i.e. subpopulation of neurons whose firing rate increases) also gets squeezed, whereas the log-rates of the subpopulation that is shifted to the left gets stretched. Both the stretch and the squeeze are signatures of a larger relative impact of the brain state change on the rates of the slow-firing neurons within each subpopulation. The combined effect of the shift and change in width of the log-rate distribution of each subpopulation produces the stretching or squeezing of the log-rate distribution of the entire neuronal population. These findings reveal the generic pattern of reorganisation of firing rates across a neuronal population as a result of different kinds of brain state transitions.

## Results

### Oral administration of TCB-2 to mice

TCB-2 is a high affinity, potent 5-HT2AR agonist (McLean et al., 2006; Fox et al., 2010). We chose to use it for our study since it is highly specific to this receptor subtype (McLean et al., 2006; Di Giovanni and De Deurwaerdère, 2018). In rodents, psychedelic drugs are typically administered by injection. Here, instead, we opted to develop an oral administration protocol, as we reasoned that oral delivery would more closely model oral administration in human subjects and would be less stressful for the animals, thus minimising any confounding neuronal activity elicited by the drug administration rather than by the drug itself. Since TCB-2 has a bitter taste, we gradually habituated the animals to the drinking of a bitter-sweet liquid, containing sucrose and a(n inert) bitterant. The concentration of the bitterant was gradually increased over the course of several days of the habituation. This allowed us to deliver the TCB-2 drug orally, dissolved in a sucrose solution, voluntarily consumed by the animals (see Methods for further details).

To confirm the effectiveness of oral administration, we tested the animals for the presence of head twitch responses (HTRs) – a motor response which in rodents is highly specific to psychedelic compounds (Willins and Meltzer, 1997; Canal and Morgan, 2012; Halberstadt and Geyer, 2013). In each test session, an observer scored the number of HTRs of two mice, one of which received sucrose liquid with TCB-2 while the other received a liquid with a bitterant but without the drug. The observer was blind to the content of the liquid each mouse received. Across the 6 test sessions, there was a consistent and pronounced increase in the number of HTRs in the mouse that received the drug (Figure 1A). We conclude that our oral delivery protocol is an effective method for systemic administration of TCB-2.

**Figure 1.**
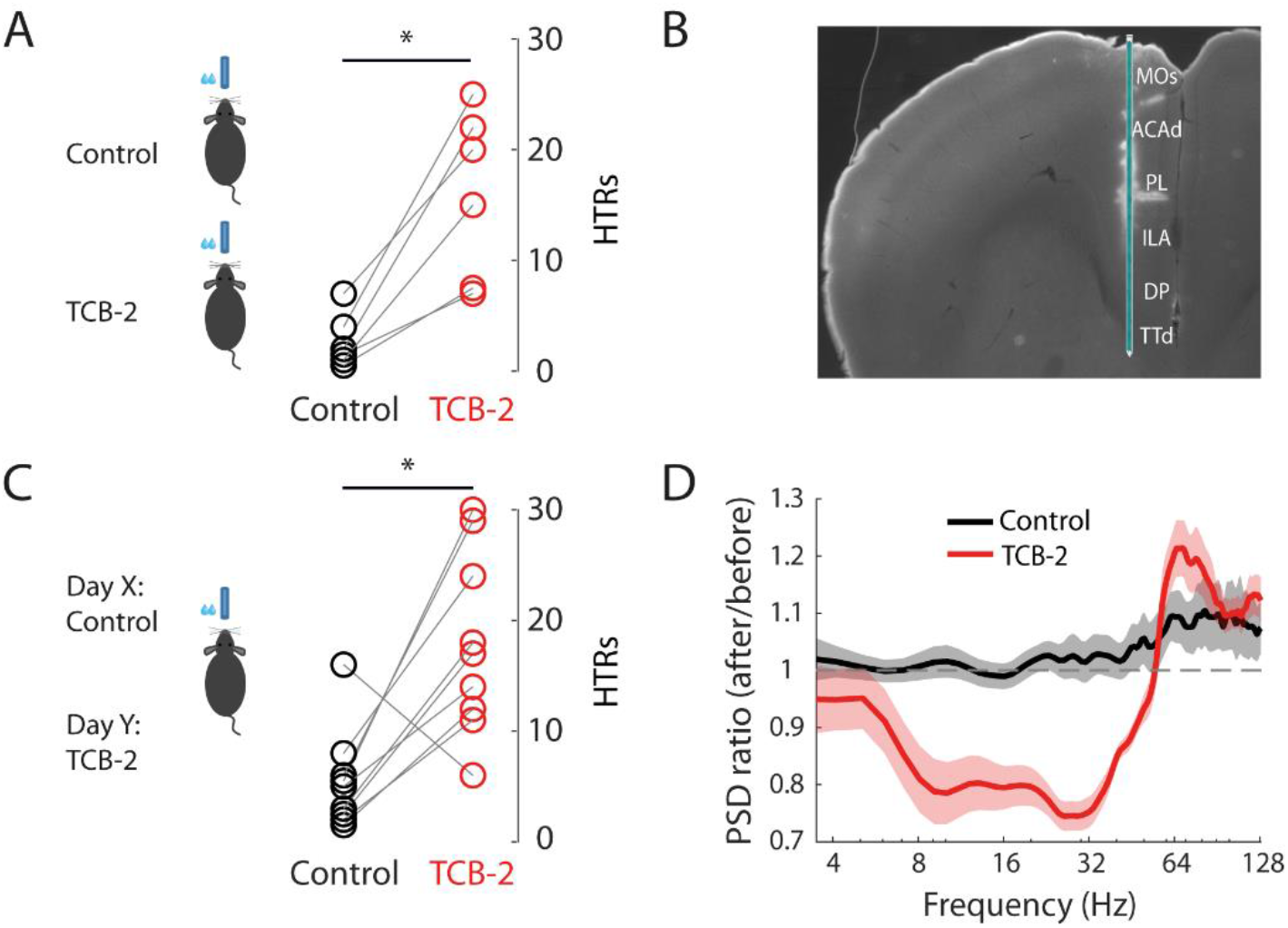
Oral administration of TCB-2 elicits HTRs and a reduction in low frequency LFP power in mPFC. **(A)** Number of HTRs observed during a 9 minute interval in 6 pairs of freely moving animals in which one animal received TCB-2 while the other received a control bitterant liquid (P = 0.016, binomial test). **(B)** Illustration of the placement of Neuropixels probe in the mPFC during the recordings (MOs – secondary motor cortex, ACAd – dorsal anterior cingulate area, PL – prelimbic cortex, ILA – infralimbic area, DP – dorsal peduncular area, TTd – dorsal taenia tecta), based on histological reconstruction of the probe track. **(C)** Number of HTRs observed during a 9 minute interval in 9 mice at the end of the recordings (one control and one TCB-2 recording per animal, P = 0.020, binomial test). **(D)** Ratio between the baseline LFP power and LFP power in the end of the recording (in each case estimated from a 10 minute interval) in control (black) and TCB-2 (red) conditions (mean of 13 recordings, shaded area shows SEM).

To record field potential and spiking activity in the medial prefrontal cortex (mPFC, Figure 1B), a new batch of mice were implanted with a metal headplate under general anaesthesia (see Methods). After several days of recovery, the animals were acclimatised to head fixation and to the drinking of bitter-sweet liquid while head-fixed. On separate days, we performed two recordings in each animal, in one of which the animal received the drug and in the other the control bitter-sweet liquid. Each recording lasted 80-120 minutes, starting with ≥ 20 minutes of baseline activity, followed by 15-20 minutes of drinking and a post-drinking interval of ≥ 30 minutes. The order of the control and drug recordings was decided randomly for each animal, and the experimenter was blinded to the treatment. In 9 of the 14 recorded animals, HTRs were observed at the end of each recording session, again showing a clear distinction between the two conditions within subjects, at a longer interval after administration (Figure 1C, Supplementary video). Consistent with previous reports on the effect of psychedelic drugs on local field potential (LFP) in the cortex (Wood et al., 2012; Michaiel et al., 2019; Thomas et al., 2021), we found that TCB-2 reduced the LFP power in the beta and low gamma frequency range by ~20%, with a smaller but significant increase in high gamma LFP power (Figure 1D).

### Temporal dynamics and magnitude of TCB-2 driven changes in single neuron activity

We next examined the temporal dynamics and the magnitude of changes in the firing of individual mPFC neurons caused by TCB-2. The firing rate of cortical neurons of awake animals is highly variable across all timescales (McCormick et al., 2015; Musall et al., 2019; Okun et al., 2019; Stringer et al., 2019), thus in order to quantify the effect of TCB-2, we fitted the firing of each neuron over an interval which included 19.5 minutes pre-drinking and 48.5 minutes post-drinking onset with a step function. This fit provided the approximate point in time at which the most prominent change in the neuron’s firing rate occurred, and the magnitude of this change (Figure 2A, see Methods). Because of firing variability, the fit finds a change point even in neurons with small or no change in firing rate; thus, for the analysis of the timing of the firing rate changes caused by the drug we considered only units in which the step function fit indicated > 2-fold change (quantitatively similar results were obtained with other thresholds, e.g., 1.5 or 2.5). We found that in both control and TCB-2 experiments, change points clustered around the time point at which the spout was made available to the mouse (resulting in a prominent change in behaviour). In addition, in TCB-2 recordings, but not in control recordings, we observed a second prominent cluster of change points, with a peak ~12 minutes after drinking onset (Figure 2B-D). Correspondingly, the percentage of units with > 2-fold change in firing rate whose change point fell between 3 and 18 minutes after the start of drinking (the temporal location of this second cluster) was significantly higher in TCB-2 recordings compared to the controls, both for units that increased their firing rate (10.8% vs 5.7%, Figure 2E) and for units that decreased their firing rate (21.1% vs 4.9%, Figure 2F). These findings indicate that within minutes from the beginning of its administration, TCB-2 produced marked changes in neuronal firing.

**Figure 2.**
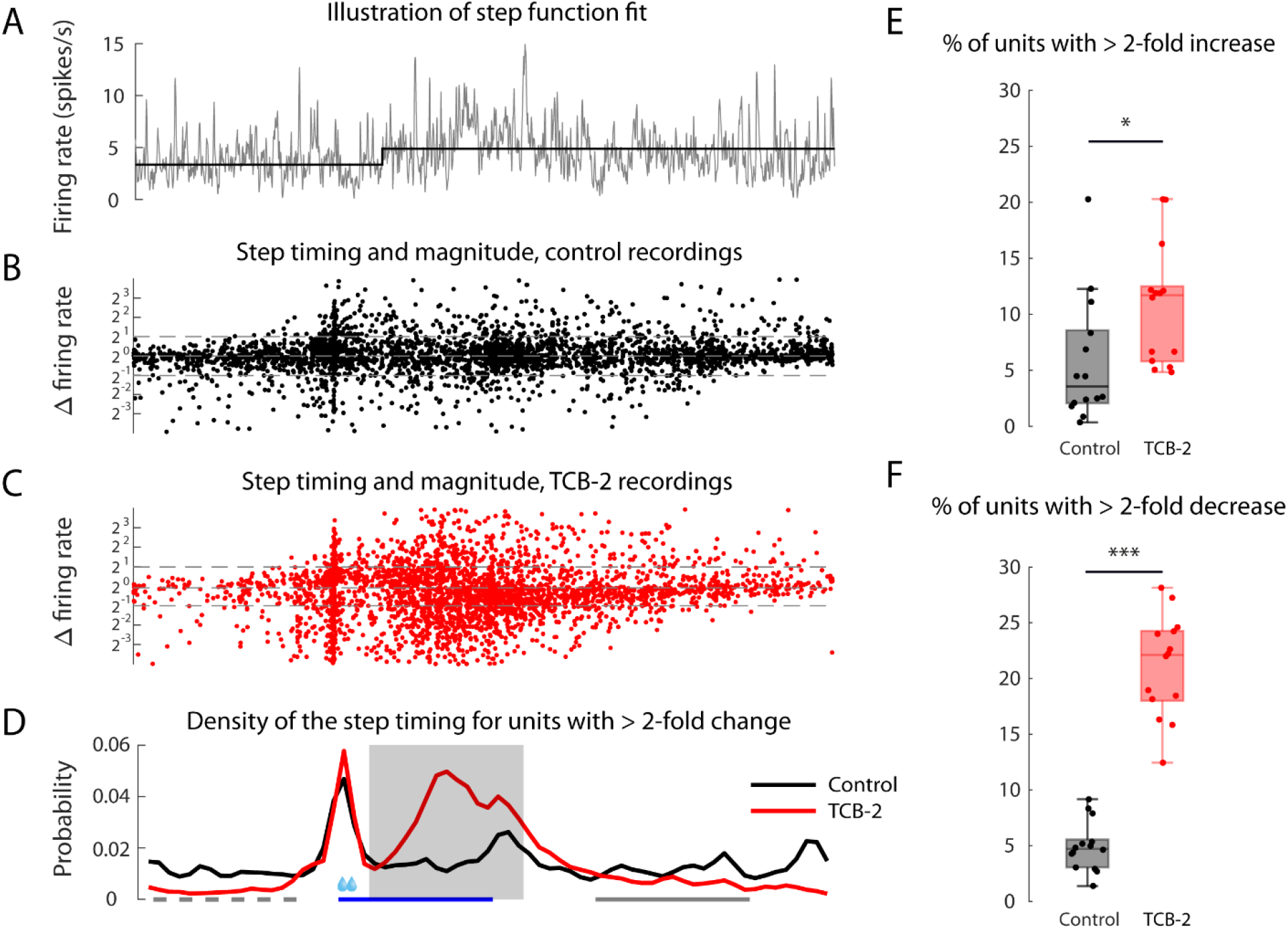
Dynamics of changes in firing rate of mPFC neurons following oral administration of TCB-2. **(A)** Illustration of step function fitting to the highly variable firing of an example well-isolated unit. **(B, C)** The position in time (x-axis) and the relative magnitude (y-axis) of the change point for each unit in the control and TCB-2 recordings, correspondingly (40/3612 and 335/3564 units are outside the y-axis limits and hence are not visible in B and C, correspondingly; 14 recordings in each condition). **(D)** Density of change point timing of units with > 2-fold change in firing rate in control and TCB-2 recordings. Blue bar indicates the 15-minute drinking interval, starting from the point at which the spout became available to the animal. **(E)** Percentage of units whose step function fit shows > 2-fold increase in firing occurring 3-18 minutes after drinking onset (this interval is indicated by the shaded area in D). This percentage was significantly higher in TCB-2 recordings (means of 10.8% vs 5.7%, P = 1.5×10^−3^, t-test; 14 recordings in each condition). **(F)** Same format as (E) for units whose step function fit shows > 2-fold decrease occurring 3-18 minutes after drinking onset. In TCB-2 recordings, the percentage of such units was significantly higher (21.1% vs 4.9%, P = 1.2×10^−7^, 14 recordings in each condition). Grey (dashed and continuous) bars in (D) indicate the ‘before’ and ‘after’ intervals used in subsequent analyses.

To characterise the reorganisation of firing rates caused by the drug, we considered the distribution of firing rates before and after drinking in the control and TCB-2 conditions. For this analysis we did not rely on the step function fit, and simply used the mean firing rate of every neuron in a 15-minute interval before drinking started and in a 15-minute interval starting 25 minutes after drinking onset (by which time drinking had stopped and the drug effect fully kicked in, as seen in Figure 2D). In the control recordings, firing rates in the majority of neurons were stable across these ‘before’ and ‘after’ intervals, i.e., most of the weight of the distribution abutted the diagonal (Figure 3A-B). In contrast, in TCB-2 recordings this distribution was much wider, indicating that the drug produced a more substantial change in firing rates of many neurons (Figure 3C-E). To quantify the magnitude of this effect, for each unit we computed the absolute rate modulation index, defined as |log(*r*_*a*_/*r*_*b*_)|, where *r*_*b*_ and *r*_*a*_ are the ‘before’ and ‘after’ firing rates of the unit (here and elsewhere log denotes a base 10 logarithm). The values of the absolute rate modulation index were significantly higher in TCB-2 recordings, when compared to the controls (Figure 3F). This was also the case when units whose firing rate increased and units whose firing rate decreased were considered separately (medians of 0.12 vs 0.24, P = 3.6×10^−45^; and 0.10 vs 0.33, P = 2.8×10^−129^, respectively; rank-sum test). All the differences remained highly significant also at the level of individual animals (at P < 10^−3^ the medians for units with increasing rates were significantly different in 9 of the 14 animals, and for units with decreasing rates this was the case in 10 of the 14 animals).

**Figure 3.**
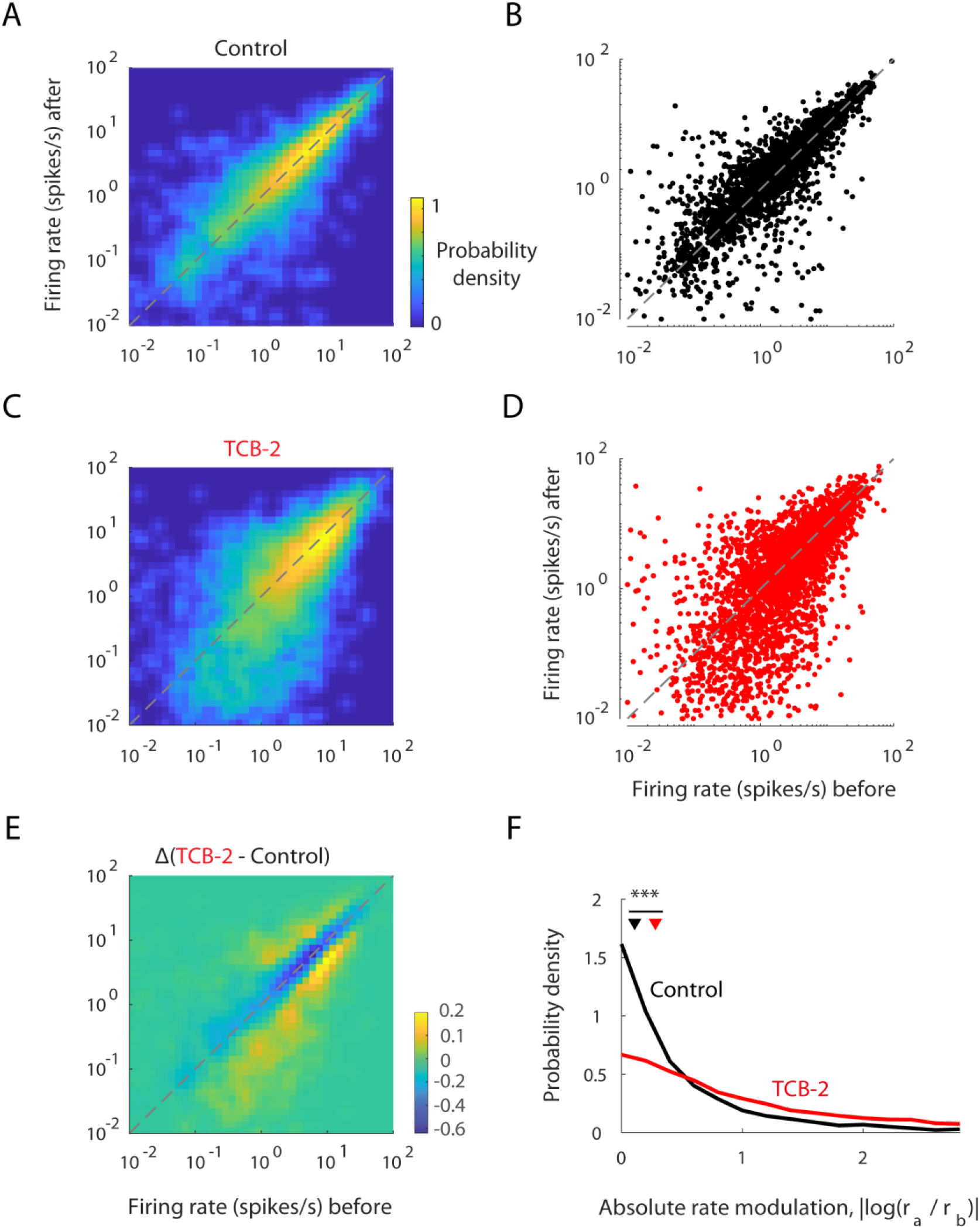
Changes in firing rates in mPFC following administration of TCB-2. **(A, B)** Distribution of firing rates before and after drinking in the control condition. In (A) the distribution is represented as a colour coded two-dimensional density plot, whereas in (B) the underlying data points corresponding to individual neurons are shown (3612 units, 14 recordings, 58 units not seen). **(C, D)** same format as (A, B) for TCB-2 condition (3564 units, 14 recordings, 160 units not seen). **(E)** The difference between the distributions shown in (C) and (A). Compared to the control condition, the TCB-2 condition had an excess (yellow) of neurons whose firing rate increased or decreased, and a lower proportion of neurons with a stable firing rate (blue). **(F)** Absolute rate modulation index distribution for units in the control and TCB-2 conditions. Medians of the two distributions are indicated by ∇ (0.11 and 0.28, correspondingly, P = 3.4×10^−174^, rank-sum test).

### TCB-2 stretches the distribution of log firing rates

Does administration of TCB-2 result in a population level change in the firing rates? Since the changes in firing rate were not unidirectional, but rather some neurons’ firing rate was elevated while others’ firing rate was suppressed, the overall distribution might not have been affected. However, visual inspection of the cumulative distributions of the firing rates suggested that the distribution after ingesting the drug was prominently different from the distribution after drinking the control liquid (Figure 4A). Indeed, in the TCB-2 condition, the mean log-rate after ingesting TCB-2 was significantly lower than in the baseline (before: 0.36 vs after: 0.25, P = 2.2×10^−10^; t-test), whereas after ingesting the control bitterant this was not the case (before: 0.34 vs after: 0.37, P = 0.13). However, an inspection of the probability density function of the log-rate distribution, obtained through analytical fitting (see Methods, Figure S1A), indicated that the primary effect of TCB-2 was not shifting the distribution to the left, as one might have expected based on the comparison of the means, but rather stretching it (Figure 4B). Indeed, in the TCB-2 condition, the standard deviation of the log-rates after drug ingestion was significantly higher than in the baseline (before: 0.64 vs after: 0.80, P = 6.4×10^−35^, Levene’s test), whereas in the control condition there was no significant change (before: 0.68 vs after: 0.67, P = 0.42).

**Figure 4.**
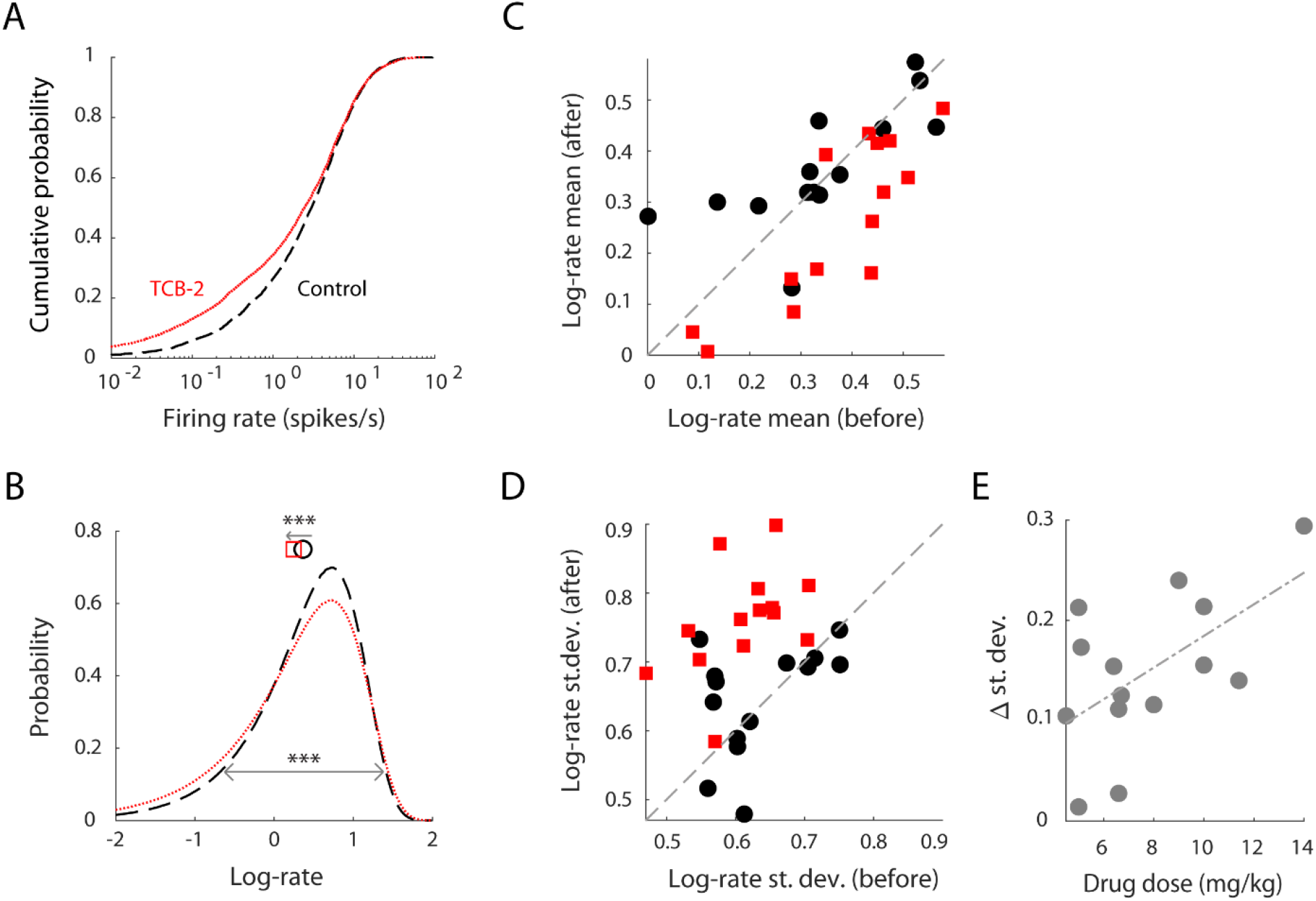
TCB-2 stretches and lowers the mean of the log firing rate distribution in mPFC. **(A, B)** Cumulative distribution of neuronal firing rates after administration of control (black, dashed) and TCB-2 (red, dotted) liquids and the probability density of a log-gamma distribution fit to the log-rates (3612 neurons from 14 recordings in the control condition, 3564 neurons from 14 recordings in the TCB-2 condition). In (B) the means of the control and TCB-2 empirical distributions are indicated by ○ and □, respectively. In the TCB-2 condition the log-rates mean was significantly lower than in the control condition (0.25 vs 0.37, P = 3.1×10^−11^, t-test) and the standard deviation was significantly higher (0.80 vs 0.67, P = 3.2×10^−23^, Levene’s test). **(C, D)** Mean (in C) and standard deviation (in D) of log-rates before and after drinking in control and TCB-2 conditions, in the individual recordings (256±138 well isolated units per recording). In the control condition, no significant changes in the mean or standard deviation were observed (P = 0.34, 0.55, correspondingly; t-test, 14 recordings). In TCB-2 recordings, the means decreased and the standard deviations increased (P = 2.2×10^−10^, 6.4×10^−35^, respectively; t-test, 14 recordings). **(E)** Increase in standard deviation of the log-rate distribution was significantly correlated with the dose of TCB-2 consumed by each mouse (r = 0.58, P = 0.015, 14 recordings; Pearson correlation).

We further confirmed that TCB-2 changes the mean and variance of the log-rates by analysing each recording separately. Within each recording, the mean and standard deviation of the log firing rates were calculated before and after drinking, which was possible owing to the high yield of the Neuropixels probes, typically providing several hundred well isolated units per recording. In control recordings, the mean log-rates before and after drinking were not significantly different (Figure 4C). Similarly, there was no significant change in the standard deviation of the log-rates (Figure 4D). On the other hand, in the TCB-2 condition the means of the log-rates were significantly lower and the standard deviations were significantly higher after drinking (Figure 4C-D).

Remarkably, the prominent change in log-rates was not accompanied by any effect on the average of the firing rates in control or TCB-2 conditions (5.2 vs 5.3 spikes/s, P = 0.17 in control recordings; 5.1 vs 5.0 spikes/s, P = 0.75 in TCB-2 recordings; t-test on the means of 14 individual recordings). In other words, TCB-2 had no effect on the total number of spikes emitted by the neuronal population as a whole, rather it changed the manner in which these spikes were distributed across cells in the neuronal population. Mathematically this can be easily understood if one remembers that for a random variable *X* > 0, *mean*(*X*) depends not only on *mean*(log(*X*)) but also on *var*(log(*X*)), and a decrease of the former can be counterbalanced by an increase of the latter.

Since TCB-2 was administered through a licking spout providing the drug, the dose that each mouse received varied depending on the consumed volume of the liquid and the weight of the animal. This allowed us to ask whether the change in the width of the distribution of the log-rates correlated with the ingested dose of the drug. A significant positive correlation between the two was indeed found (r = 0.58, Figure 4E).

### Stretching and squeezing of the log-rate distribution by changes in arousal level

In the literature, modulation of the width of the firing log-rates distribution was reported for transitions between wakefulness and non-REM and REM sleep (Watson et al., 2016; Miyawaki et al., 2019; Brecier et al., 2021; Niethard et al., 2021). Arguably, a transition into a psychedelic brain state is not equivalent to falling asleep or waking up. Is it then possible that stretching (or squeezing) of log-rates is a general phenomenon that accompanies different kinds of brain state transitions? To address this question, we turned to examine pupil-indexed changes in arousal – a prominent, intrinsic and naturally occurring modulation of brain state in awake subjects (Polack et al., 2013; Reimer et al., 2014; McCormick et al., 2015; Vinck et al., 2015; Larsen and Waters, 2018). We compared the distribution of log-rates in a state of high arousal, which for the purpose of the present analysis was defined as 25% of the recording time with the largest pupil size, to the distribution in a state of low arousal, defined as 25% of the recording time with the lowest pupil size (Figure 5A, Figure S2). This comparison was performed in the baseline interval preceding drinking (in 15 recordings from 8 animals where pupil videos were recorded). We found that increased arousal significantly stretched the log-rate distribution in mPFC (Figure 5B, Figure S1B-C).

**Figure 5.**
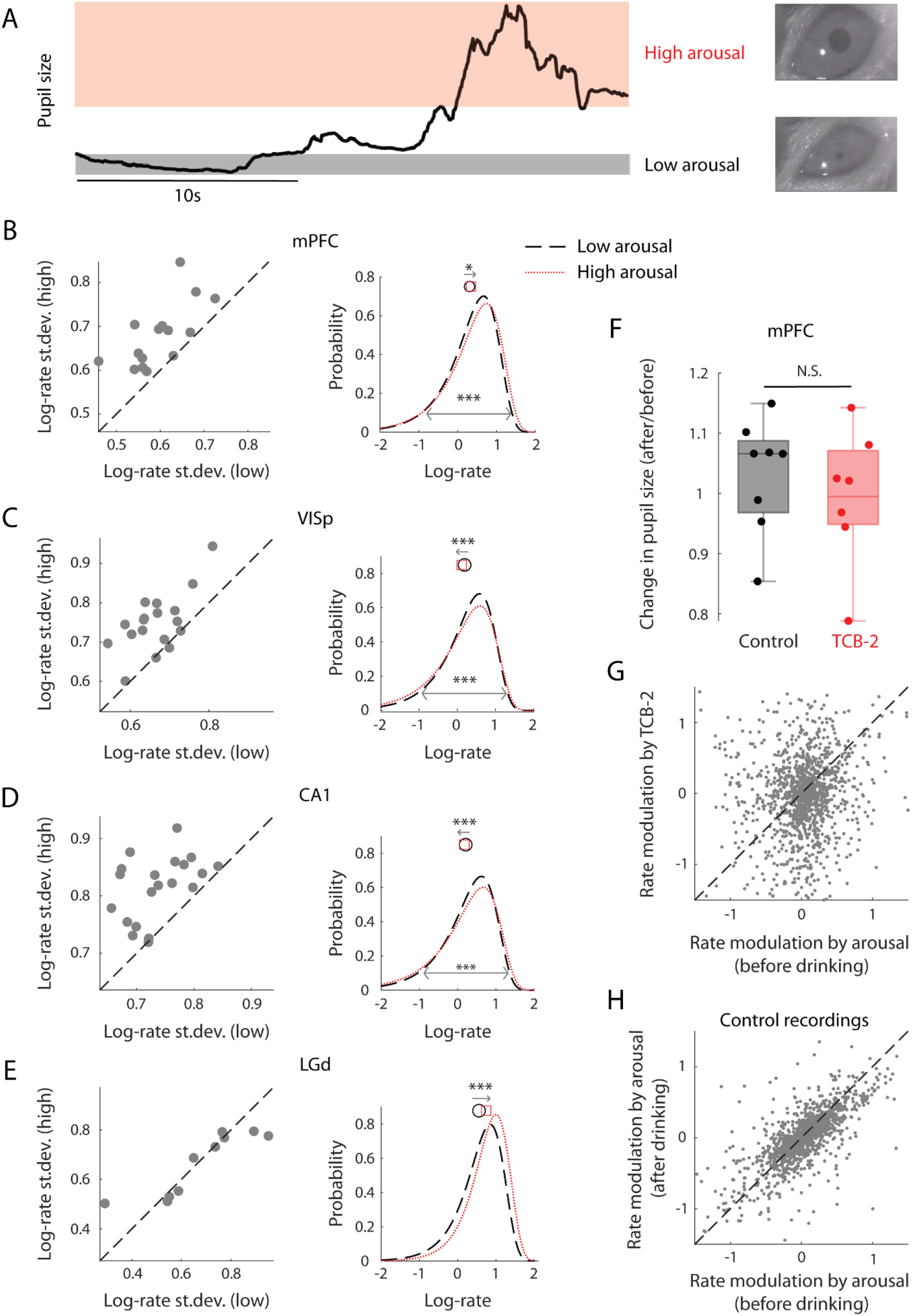
Arousal stretches the neuronal log-rate distribution in the cortex, but not in the thalamus, and does not explain the psychedelic firing rate modulation. **(A)** Illustration of the analysis: 25% of the recording interval during which pupil size was the smallest (largest) were designated as the low (high) arousal brain state. **(B)** Left: Standard deviation of the log-rates in low and high arousal brain states (15 mPFC recordings in 8 animals, 207±123 units per recording). The standard deviation was significantly higher in the high arousal state (P = 5.8×10^−5^, t-test). The standard deviation was also significantly higher in the pooled distribution (3100 neurons from 15 recordings, combined; 0.71 vs 0.64, P = 5.7×10^−8^, Levene’s test). Right: log-gamma fit of the distribution of log firing rates in low (dashed, black) and high (dotted, red) arousal states, illustrating the stretching. The means of the low and high arousal empirical distributions are indicated by ○ and □, respectively; they were also significantly different: P = 0.027, t-test. **(C)** Same format as (B), for recordings from VISp (left: P = 1.9×10^−5^ for comparison of st.dev. in individual recordings using t-test; right: P = 5.3×10^−11^ for comparison of variances in the pooled distribution using Levene’s test, P = 5.1×10^−5^ for comparison of means in the pooled distribution using t-test; 2831 neurons from 18 recordings). **(D)** Same format as (C) for recordings from CA1 (left: P = 7.0×10^−6^, right: P = 5.7×10^−21^ for comparison of variances, P = 9.1×10^−4^ for comparison of means; 7729 neurons from 20 recordings). **(E)** Same format as (C) for recordings from LGd (left: P = 0.76, right: P = 0.63 for comparison of variances, P = 1.4×10^−7^ for comparison of means; 690 neurons from 10 recordings). **(F)** TCB-2 administration did not produce any significant change in pupil size (P = 0.53, t-test of 8 control recordings vs 7 TCB-2 recordings). Furthermore, pupil area did not significantly change during either control or TCB-2 recordings (control: P = 0.31, TCB-2: P = 1, signed rank test). **(G)** Across single units, the correlation between modulation by arousal and by TCB-2 was low (r = 0.11, Spearman correlation, P = 4.2×10^−5^, 1356 units from 7 recordings; 71 units are beyond the axes limits seen in the plot). **(H)** Correlation between modulation by arousal before and after drinking in recordings with control bitterant was high (r = 0.74, P = 2.1×10^−299^, 1744 units from 8 recordings; 18 units are beyond the axes limits seen in the plot).

To further test the robustness of this last finding and to examine whether it extends to other cortical and subcortical areas, we used a large scale dataset of Neuropixels recordings in the mouse that was recently made publicly available by the Allen Brain Institute (Siegle et al., 2021). In this dataset, we compared the distribution of log-rates in low and high arousal, similarly to the analysis of our own mPFC recordings. We excluded periods of running, limiting our analysis to periods of immobility, to avoid introducing a behavioural state which did not exist in our own recordings. The comparison showed that both in the primary visual cortex (VISp) and in the hippocampal area CA1, increased arousal resulted in a prominent stretching of the distribution of log-rates (Figure 5C-D, Figure S1D-G). Interestingly, this was not the case in the lateral geniculate nucleus (LGd), where arousal resulted in an increase in the mean of the distribution and a non-significant squeeze rather than a stretch (Figure 5E, Figure S1H-I). We conclude that the stretching of the log-rate distribution is not unique to the transition into the psychedelic brain state, and occurs in intrinsic, naturally occurring brain state transitions such as arousal changes.

The finding that arousal stretches the distribution of log-rates suggests the possibility that the effect of TCB-2 is explained by arousal. To test this potential explanation, we compared the pupil size before and after drinking in TCB-2 and control bitterant recordings. We found no significant difference between the two, furthermore in both cases there was no significant change in pupil size in the intervals before and after drinking (Figure 5F). This observation is in agreement with (Michaiel et al., 2019), who also observed no effect of a psychedelic compound on mouse pupil size.

Next, at the level of individual neurons, we compared the modulation of firing rates by arousal to modulation by TCB-2. Towards this end, we examined the correlation between the index of firing rate modulation by TCB-2 (log(*r*_*a*_/*r*_*b*_), where *r*_*b*_ and *r*_*a*_ are the firing rates before and after TCB-2 administration) and the index of firing rate modulation by arousal in the interval before the drinking spout was introduced (log(*r*_*h*_/*r*_*l*_), where *r*_*h*_ and *r*_*l*_ are the firing rates during high and low arousal intervals) across all the neurons. The correlation between the two indexes, while significant, was low (r = 0.11, Figure 5G; when this analysis was repeated separately within each recording, the mean correlation was 0.13±0.17, 7 recordings). For comparison, correlation between firing rate modulation by arousal in the intervals before and after drinking was 0.41 for TCB-2 recordings and 0.74 for control recordings (Figure 5H; when this analysis was repeated separately within each recording, the mean correlation was 0.47±0.19 for 7 TCB-2 recordings and 0.70±0.16 for 8 control recordings), indicating that the arousal modulation index is robust over the timescales of the recording. We conclude that the stretching of log-rates by TCB-2 was not caused by pupil-indexed arousal and that the two effects are almost orthogonal at the neuronal population level.

### Changes to the log-rate distributions of the upregulated and downregulated neuronal subpopulations underlying the modulation of the log-rate distribution of the whole population

The stretching and squeezing of the log-rate distribution either by TCB-2 or by changes in arousal level can be explained if neurons whose firing rate is elevated and neurons whose firing rate is suppressed by a transition from brain state *Y* to brain state *Z* are considered separately. In the language of probability theory, we consider the log-rate distribution across the entire neuronal population to be a mixture of two distributions, one of which is the subpopulation upregulated by the *Y* → *Z* transitions, and the other is the subpopulation downregulated by such transitions (and vice versa for *Z* → *Y* transitions). If in brain state *Y* the distributions of the upregulated and the downregulated neurons are closer to each other than in brain state *Z*, we would expect the log-rate distribution across all the neurons to be wider in brain state *Z*, as illustrated in a schematic plot in Figure 6A. Indeed, in the case of transition into the psychedelic brain state, the log-rate distributions of mPFC neurons whose firing rate was elevated (which comprised 39% of all the neurons) and of neurons whose firing was suppressed (the remaining 61%) were close to each other in the baseline condition, but after TCB-2 was ingested the two distributions were drawn apart (Figure 6B, Figure S3A). This change was highly significant when quantified by the distance between the means of the two distributions before and after drinking in individual recordings (Figure S3B). Similarly, CA1 subpopulations upregulated and downregulated by arousal were closer together in the low arousal state compared to the high arousal brain state (Figure 6C, Figure S3C-D), explaining the stretching of the overall log-rate distribution with transition into the highly aroused state (and its squeezing with transition into low aroused state). Similar dependence on arousal was also observed in VISp (Figure S3E-F) and mPFC (Figure S3G-H). Of note, for these analyses the assignment of every neuron to one of the two subpopulations and the estimation of its firing rate in the two brain states were cross-validated, by using different and non-overlapping recording segments for each task (see Methods).

**Figure 6.**
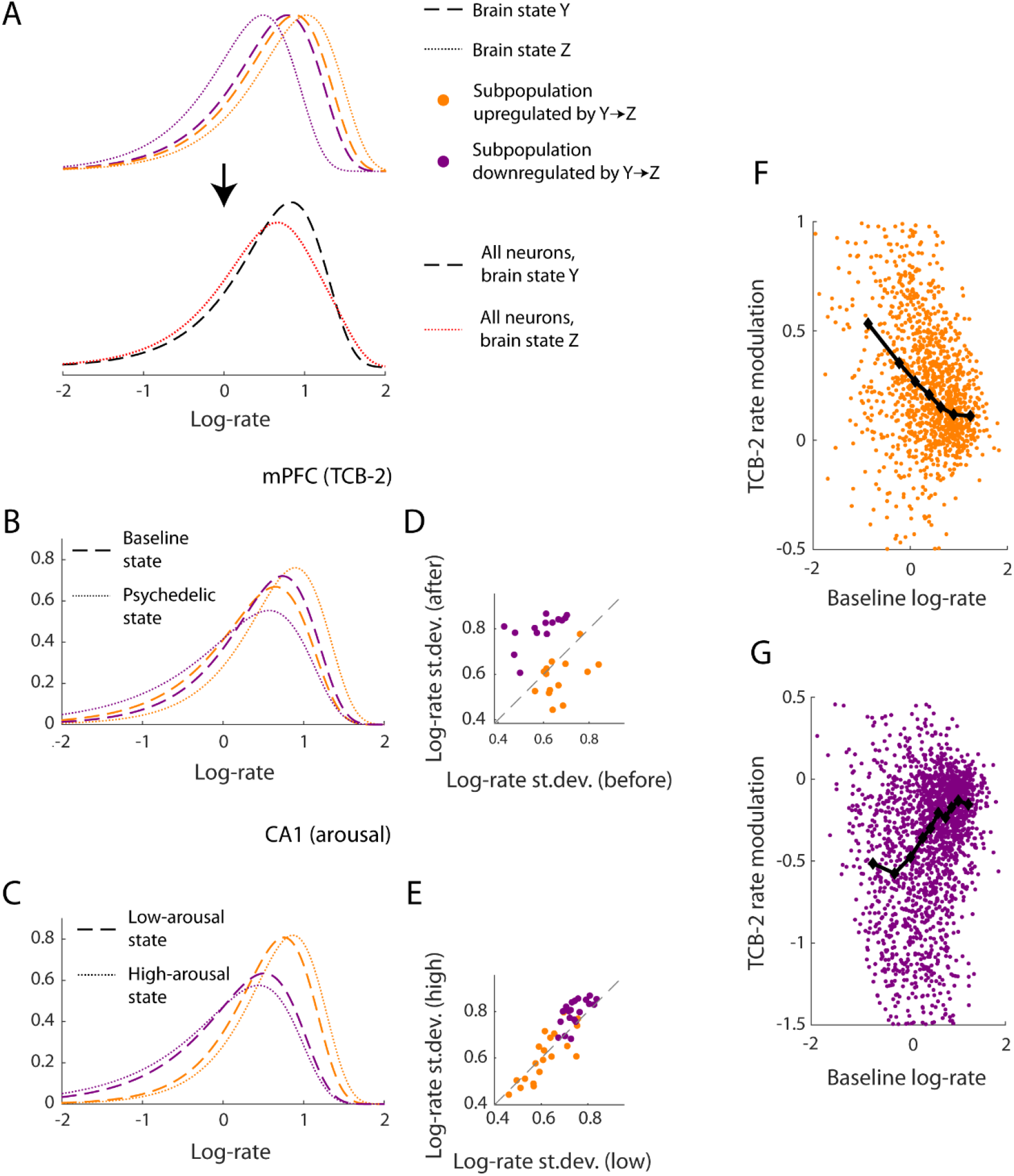
Changes in the firing rates of the upregulated and downregulated subpopulations underlie the stretching or squeezing of the log-rate distribution of the entire neuronal population. **(A)** Schematic illustration. Top: the transition from brain state *Y* (dashed lines) to brain state *Z* (dotted lines) elevates the firing rates of 50% of the neurons (orange) while suppressing the firing rates of the other 50% (purple). The two subpopulations have closely overlapping log-rate distributions in brain state *Y*, but not in brain state *Z*. Bottom: the log-rate distribution of the entire population is a combination (mixture) of the distributions of the two subpopulations. This distribution gets stretched by *Y* → *Z* brain state transitions and it gets squeezed when the brain state switches in the opposite direction. **(B)** Log-gamma fit of the distribution of log-rates of mPFC neurons whose firing rate was elevated by TCB-2 (39%, orange) and of neurons whose firing rate was suppressed by TCB-2 (61%, purple) in the baseline part of the recording (dashed line) and after drug ingestion (dotted line). The log-gamma fit of the distribution of log-rates of the entire population of mPFC neurons (the combination of the distributions of the elevated and suppressed neurons) was presented in Figure 4B. **(C)** Log-gamma fit of the distribution of log-rates of CA1 neurons whose firing rate was elevated by increased arousal (40%, orange) and of neurons whose firing rate was suppressed by increased arousal (60%, purple) in low (dashed line) and high (dotted line) arousal brain states. The log-gamma fit of the distribution of log-rates of the entire population of CA1 neurons was presented in Figure 5D. **(D)** Standard deviation of the log-rates in baseline and psychedelic brain states of subpopulations of upregulated (orange) and downregulated (purple) by TCB-2. In the former subpopulation, TCB-2 ingestion led to a decrease in the standard deviation of the log-rates (0.67±0.08 vs 0.59±0.09, P = 3.3×10^−3^, t-test of the st.dev. before and after drinking, 14 recordings). In the latter subpopulation, TCB-2 ingestion led to an increase in the standard deviation of the log-rates (0.59±0.09 vs 0.80±0.07, P = 5.7×10^−8^, t-test of the st.dev. before and after drinking). **(E)** Standard deviation of the log-rates in low- and high-arousal brain states of CA1 subpopulations upregulated (orange) and downregulated (purple) by an increase in arousal. In the former subpopulation, changes in arousal did not change the st.dev. of the log-rate distribution (0.64±0.08 vs 0.63±0.11, P = 0.26, t-test, 20 recordings). In the latter subpopulation, high arousal led to an increase of st.dev. of the log-rate distribution (0.75±0.05 vs 0.80±0.06, P = 7.7×10^−5^, t-test). **(F)** In the mPFC subpopulation of neurons upregulated by TCB-2, the modulation of firing rate caused by the drug had a significant negative correlation with their baseline log-rate (r = −0.42, P = 4.8×10^−59^, 1388 units, Spearman correlation). Black diamonds – running median. A small proportion of units classified as upregulated by TCB-2 had a negative rate modulation index; this is inherent to cross-validation between assignment of units to the two subpopulations and estimation of their firing rate in the baseline and TCB-2 conditions (see Methods). **(G)** In the mPFC subpopulation of neurons downregulated by TCB-2, the modulation of firing rate caused by the drug had a significant positive correlation with their baseline log-rate (r = 0.24, P = 1.2×10^−27^, 2176 units). Black diamonds – running median. As in (F), a small proportion of units classified as downregulated had a positive rate modulation index.

Visual inspection of the log-gamma probability density function fits in Figure 6B-C suggest that the theoretical scheme depicted in Figure 6A is too simplistic in comparison to the empirical data. The theoretical scheme makes a simplifying assumption that the variance of each subpopulation is not affected by the brain state transition, whereas the fits to actual data suggest that rightward shift of a subpopulation’s log-rate distribution was accompanied by a squeeze (the peak of the distribution gets higher), whereas leftward shift was accompanied by a stretch (the peak of the distribution gets lower). Levene’s test of variance equality confirmed this suggestion for both the upregulated and downregulated subpopulations in the TCB-2 case (standard deviation of 0.69 vs 0.61, P = 9.8×10^−5^ for the upregulated subpopulation; 0.64 vs 0.85, P = 2.3×10^−51^ for the downregulated subpopulation). This was further confirmed by comparing the standard deviation of each subpopulation separately within every recording before and after ingesting the drug (Figure 6D).

The observation that a squeeze accompanies rightward shift and stretch accompanies leftward shift also held for the case of transitions in arousal level. Specifically, Levene’s test of log-rates of CA1 neurons that were downregulated by increased arousal showed that their variance was significantly higher in the high arousal brain state (standard deviations of 0.82 vs 0.75, P = 8.7×10^−10^). We further confirmed this by comparing the standard deviation of the log-rates of this subpopulation between high and low arousal conditions within individual recordings (Figure 6E). The upregulated CA1 subpopulation showed no significant change in variance between the high and low arousal brain states (P = 0.92 Levene’s test on all the units, P = 0.80 when comparing standard deviations across recordings, Figure 6E). Similarly, modulation of VISp and mPFC by arousal stretched or squeezed the log firing rate distribution of neurons which were downregulated by increase in arousal but had no significant effect on the variance of the distribution of the subpopulation upregulated by increase in arousal (Figure S4). In LGd, the proportion of neurons downregulated by increased arousal was low (just 14%), thus the mean and variance of log-rates of the entire neuronal population were determined by neurons whose rate was upregulated by increased arousal. This explains why LGd was the only examined brain area where the primary effect of increased arousal on the log-rate distribution of the entire population was a rightward shift rather than a stretch (Figure 5).

A squeeze of the log-rate distribution of a neuronal subpopulation that is upregulated by a brain state transition implies the existence of a correlation between the firing rate and the modulation magnitude within this subpopulation. To demonstrate this point, let us use *r*_*i*_ to denote the firing log-rate of neuron *N* ≥ *i* ≥ 1 in brain state *Y* (with lower firing rates), and let *m*_*i*_ + *r*_*i*_ be its log-rate in brain state *Z* (with higher firing rates), where *m*_*i*_ is the firing rate modulation index. In other words, the firing rate of neuron *i* changes from 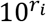 to 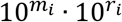, which is the same simple multiplicative model of modulation used in our previous analyses. Let *R* and *M* denote the distributions of the log-rates and their modulation indexes across the entire subpopulation of neurons. It holds that *Var*(*R* + *M*) = *Var*(*R*) + *Var*(*M*) + 2 ∙ *Cov*(*R*, *M*). Because the overall effect of the modulation is a squeeze of the distribution, i.e., *Var*(*R* + *M*) < *Var*(*R*), we can conclude that *Cov*(*R*, *M*) < −*Var*(*M*)/2 ≤ 0. Thus, we find that the modulation index is negatively correlated with the original log-rate, i.e., the modulation has to be larger in neurons with lower firing rates.

Next, we tested this mathematical inference for the psychedelic brain state transition. Within the mPFC subpopulation of neurons upregulated by TCB-2, we found a substantial negative correlation between the baseline log-rate of the neurons and their firing rate modulation by the drug (Figure 6F). It is noteworthy that on one hand this negative correlation is predicted by the squeeze, as was argued above, but on the other hand it also explains the squeeze. Indeed, given that as a result of TCB-2 administration the log-rates of the upregulated fast-firing neurons shift to the right less than the log-rates of the upregulated slow-firing neurons, the overall log-rate distribution of the upregulated neurons must get squeezed. Within the subpopulation of neurons downregulated by TCB-2 we observed the reverse effect: there was a substantial positive correlation between the baseline log-rate of the neurons and their firing rate modulation by the drug (Figure 6G). The negative correlation in the upregulated subpopulation and the positive correlation in the downregulated population combine to produce a significant negative correlation between the absolute magnitude of rate change produced by TCB-2 (the absolute rate modulation index) and the baseline log-rate, when the entire neuronal population was considered (r = −0.39, P = 8.0×10^−121^, 3564 units from 14 recordings in mPFC, Spearman correlation). This was also seen at the level of individual recordings (mean correlation = −0.36±0.06, P< 0.01 in 14/14 recordings, Spearman correlation).

Finally, we tested the relationship between neuronal firing rate and magnitude of its modulation by brain state transitions across different levels of arousal. Similar to the case of TCB-2, the subpopulations upregulated by increased arousal in mPFC, VISp, CA1 and LGd had negative correlation between log-rates (in the low arousal state) and the arousal modulation index (r = −0.28, P = 3.0×10^−33^, 1828 units from 15 recordings in mPFC; r = −0.17, P = 2.0×10^−8^, 1149 units from 18 recordings in VISp; r = −0.11, P = 1.7× 10^−8^, 2979 units from 20 recordings in CA1; r = −0.46, P = 4.5×10^−30^, 580 units from 10 recordings in LGd, Spearman correlation). Conversely, the mPFC, VISp and CA1 subpopulations downregulated by increased arousal exhibited a positive correlation between log-rates and the arousal modulation index (r = 0.25, P = 4.0×10^−19^, 1272 units from 15 recordings in mPFC; r = 0.15, P = 1.5×10^−9^, 1682 units from 18 recordings in VISp and r = 0.30, P = 6.1×10^−88^, 4750 units from 20 recordings in CA1, Spearman correlation). The negative correlation in the subpopulation upregulated by increased arousal and the positive correlation in the downregulated population combine to produce a significant negative correlation between the absolute magnitude of rate change produced by arousal and the log-rate (in low arousal), when the entire neuronal population was considered (r = −0.42, P = 2.4×10^−130^, 3100 units from 15 recordings in mPFC; r = −0.36, P = 8.0×10^−81^, 2831units from 18 recordings in VISp; r = −0.43, P = 3.4×10^−319^, 7729 units from 20 recordings in CA1; r = −0.46, P = 4.0×10^−36^, 690 units from 10 recordings in LGd, Spearman correlation). The correlation between absolute rate modulation and low arousal log-rates of all the neurons was also seen at the level of individual recordings: mean correlation across all recordings = −0.41±0.09 with P < 0.01 in 15/15 animals in mPFC; −0.35±0.12 with P < 0.01 in 17/18 animals in VISp; −0.42±0.07 with P < 0.01 in 20/20 animals in CA1; −0.39±0.22 with P < 0.01 in 5/10 animals in LGd.

## Discussion

In this study, our initial goal was to characterise how systemic administration of a psychedelic compound changes the firing of neurons in the medial prefrontal cortex. We found that starting several minutes after the beginning of oral delivery of the TCB-2 drug, the firing rates were bidirectionally modulated, with 39% and 61% neurons up- and down-regulated, respectively (Figure 2, Figure 3). Our key observation was that as a result of this modulation, the distribution of the firing log-rates of a neuronal population was prominently stretched (Figure 4, log-rates were used throughout our analyses because the natural scale for the distribution of firing rates is logarithmic, e.g., see Hromadka et al., 2008; Roxin et al., 2011; Buzsaki and Mizuseki, 2014). Subsequently, we investigated whether stretching or the reverse effect of squeezing are typical in other brain state transitions. We found that naturally occurring transitions into a more (less) aroused brain state, as indicated by pupil dilation, also resulted in the stretching (squeezing) of the log-rate distribution in the cortex and hippocampus. Arousal-driven stretching did not, however, explain the psychedelic effect, since pupil size was not significantly different in the ‘tripping’ mice, and at the level of individual neurons the arousal and psychedelic modulation indexes were only weakly correlated, i.e., the two classes of brain state modulation were almost orthogonal (Figure 5G). Thus, stretching and squeezing of the log-rate distribution occurs in different kinds of transitions in brain state. Next, we demonstrated that such stretching or squeezing are a façade hiding an interplay between the log-rate distributions of the up- and down-regulated subpopulations of neurons (Figure 6). Specifically, the change in the log-rate distribution across the entire neuronal population is determined by the changes in the mean and variance of each subpopulation, and the size of each subpopulation relative to the other. Finally, we observed that across all the examined brain state transitions and areas, a squeeze of the log-rates distribution of a subpopulation of unidirectionally modulated neurons occurs together with an increase in firing rates and a reverse effect of a stretch occurs together with a decrease in firing rates. In both cases this means that the fast-firing neurons tend to be relatively less modulated than the slow-firing neurons (Figure 6F-G), an observation that also explains why a subpopulation which shifts to the right gets squeezed and a subpopulation which shifts to the left gets stretched.

Stretching and squeezing of the log-rate distribution were previously observed in neuronal populations in cortical and hippocampal areas across transitions between wakefulness, non-REM and REM sleep (Watson et al., 2016; Miyawaki et al., 2019). Modulation of the width of activity level distribution between sleep and wakefulness was also recently reported in studies utilising 2-photon microscope imaging (Brecier et al., 2021; Niethard et al., 2021). Miyawaki et al. (2019) report that transitions from non-REM to REM stretched the log-rate distribution in both the cortex and the hippocampus, whereas brain state transitions in the opposite direction produced a squeeze. The authors further divided the neurons into quintiles according to the average firing rate of each cell and analysed how the brain state transition affects the firing rates within each quintile. They report that in hippocampal area CA1 during transitions between non-REM and REM states, the firing rates of neurons in the top and bottom quintiles move in the opposite directions (i.e. when the top quintile shifts right, the bottom quintile shifts left, and vice versa). This example finding and their further observations concerning other cortical areas or pairs of brain states appear to be consistent with the two subpopulations scheme proposed in the present work (Figure 6).

We are aware of two previous works that studied how systemic administration of psychedelic drugs affects the spontaneous firing rates of individual cortical neurons in the non-anaesthetised mammalian brain (Wood et al., 2012; Michaiel et al., 2019). The former work relied on recordings in the prefrontal cortex of freely moving rats, whereas the latter used recordings in the primary visual cortex of head-fixed mice. Consistently with our present findings, these earlier publications describe a bidirectional modulation of the neurons, with a preponderance of downregulated neurons. The former study also suggested that the overall effect is a decrease of population activity, which at first sight might seem inconsistent with our observation that the mean firing rate was not substantially modulated. However, this conclusion of (Wood et al., 2012) was based on an analysis that normalised (z-scored) the activity of every neuron, whereas we have shown here that the population firing rate stayed relatively constant because the decrease in the firing rate of a larger proportion of neurons after TCB-2 ingestion was counterbalanced by an increase in the firing rate of a smaller proportion of neurons with higher firing rates. Our observation therefore settles this apparent inconsistency and allows us to conclude that the experimental observations reported in the three studies are in full agreement with each other.

In the last decade a major research effort was directed at understanding the effects of arousal on ongoing and sensory-evoked neuronal activity (Harris and Thiele, 2011; Busse et al., 2017; Pfeffer et al., 2021). In the majority of these studies, carried out in mice, arousal was operationally defined through pupil size or running speed. In agreement with our present data, it was generally observed that arousal produced a bidirectional modulation of ongoing activity (Niell and Stryker, 2010; Erisken et al., 2014; Vinck et al., 2015; Pakan et al., 2016; Dipoppa et al., 2018), with considerable differences between modulation locked to pupil size in immobile animals and modulation by running (Vinck et al., 2015). Recent studies utilised a plethora of methods to elucidate the mechanisms involved in this modulation, with particular focus on neuromodulation by cholinergic, noradrenergic and serotonergic inputs (Polack et al., 2013; Reimer et al., 2016; Larsen and Waters, 2018; Cazettes et al., 2021) and the distinct roles of parvalbumin (PV), somatostatin (SST) and vasoactive intestinal peptide (VIP) expressing interneurons (Polack et al., 2013; Fu et al., 2014; Reimer et al., 2014; Pakan et al., 2016; Dipoppa et al., 2018). It seems highly likely that these interneuron classes also have distinct behaviours and roles in the modulation of cortical activity by psychedelics. For example, earlier this year it was reported that a subset of SST entorhinal cortex neurons is strongly activated by their 5-HT2ARs, leading to a profound suppression of population activity by serotonin and serotonin-releasing drugs in vitro and in anaesthetised mice (de Filippo et al., 2021).

We found that slow-firing neurons are more strongly modulated than fast-firing neurons across all brain areas and all brain state transitions examined in the present work, including transitions caused by a psychedelic drug. This observation adds to the growing list of distinctions between slow-firing and fast-firing neurons. It is also in line with the general idea that the former are more ‘plastic’ whereas the latter are more ‘rigid’, recently exemplified by the finding that when a novel environment was explored, slow-firing place cells attain higher place specificity than fast-firing cells (Grosmark and Buzsaki, 2016). One mechanism that was suggested as a potential explanation for such a difference is saturation of the strength of synaptic inputs on the fast-firing neurons (Buzsaki, 2019).

Clearly, bidirectional modulation of firing rates is a widespread phenomenon which is not limited to the few specific cases that were investigated here in detail. For example, the aforementioned study of (Wood et al., 2012) examined in a separate set of recordings the effects of amphetamine, which is not a psychedelic. The authors observed that systemic administration of amphetamine also produced a bidirectional modulation of firing rates of neuronal populations in the frontal cortex, demonstrating that such modulation pattern is not unique to the psychedelic class of psychoactive drugs. Furthermore, bidirectional modulation is not limited to the brain areas examined in our work. For instance, recent population recordings in the amygdala revealed that it too exhibits a bidirectional modulation, which separates the neurons into two subpopulations that respond to social exploration either by elevation or suppression of firing. Similar (but orthogonal) bidirectional modulation was observed in the same neuronal populations during object and spatial exploration (Fustiñana et al., 2021). With these and multiple other examples in mind, we believe that the framework proposed here for analysing firing rate modulation would be useful in a wide range of studies examining firing rate modulation at the population level. Such future work would further test the generality of our observation that slow-firing neurons show relatively greater modulation than fast-firing neurons.

In summary, we found that a change in the width of the distribution of the log firing rates of a neuronal population is a commonly encountered correlate of a change in brain state. The underlying cause of the stretching or the squeezing is the interaction between a shift to the right and a squeeze of the population of upregulated neurons and a shift to the left and a stretch of the population of downregulated neurons. In both subpopulations, the relative change in firing rate is greater among the slow-firing neurons. In some cases (e.g. arousal effect in LGd) almost the entire neuronal population is shifted in the same direction, and there is no prominent second subpopulation to consider.

## Methods

### Ethical approval

All experimental procedures were conducted according to the UK Animals Scientific Procedures Act (1986) under a project license granted by the Home Office, by personal licence holders.

### Animals

Experiments were conducted on adult C57BL/6J mice of both sexes. At the time of recording, the animals were aged 8-23 weeks, and their weight was 18-32 g. Mice were housed under a 12/12 h light-dark cycle with food and water available ad libitum. All training and recordings were done during the light phase of the 24 h cycle.

### Surgeries

Mice were anaesthetised with isoflurane for all surgical procedures. For headplate implantation, after the induction of anaesthesia, the head of the animal was shaved, and hair removal cream was applied and washed off. Additional short- and long-term analgesia was provided by injection of buprenorphine into the shoulder muscles (1 mg/kg, s.c.) and by injection of bupivacaine (6 mg/kg, s.c.) under the scalp. The animal was also given an injection of a nonsteroidal anti-inflammatory drug carprofen (5 mg/kg, s.c.). Viscotears gel was used to protect the animal’s eyes while it was anaesthetised. The headplate implantation was performed in aseptic conditions using a stereotaxic frame. During the surgery, the body temperature of the mouse was maintained at ~37°C using a heated pad with a closed loop temperature controller (TMP-5, Supertech Instruments). The concentration of isoflurane was maintained at ~2%. Maintenance of an adequate depth of anaesthesia was verified by testing for the absence of the pedal withdrawal reflex with a firm pinch between the toes of the mouse and by monitoring the rate and depth of respiration throughout the surgery. The shaved skin covering the skull was cut away using forceps and scissors. The skull was scraped and thoroughly cleaned with saline and hydrogen peroxide solutions and then scored to increase the surface area for cementing of the headplate. Bregma and the craniectomy site were marked using a marker for subsequent craniectomies. The headplate was positioned horizontally on the skull, in a manner that allowed easy access to the craniectomy site and attached to it using dental cement (Super-Bond, Sun Medical). The exposed skull above the future craniectomy site was covered with Kwik-Cast (WPI). Following surgery, mice were allowed to recover in a new clean cage placed on top of a heated recovery shelf for at least 1 hour, and were provided with water-soaked food pellets and water. Mice were monitored closely until there were no signs of distress. Animals received carprieve (p.o.) for 3 days following surgery.

### Habituation and behavioural training

Mice were habituated to head fixation and to the drinking spout before recordings. To acclimatise them to the drinking spout and liquid, starting from the day after the headplate implantation surgery, mice in their home cage were offered a 15% sucrose water solution from a manually held syringe. Following at least 3 days after headplate implantation surgery, animals were gradually acclimatised to head-fixation, during which they were inside a plastic tube where they could comfortably sit or stand. Over the course of approximately a week the head-fixation duration increased from ~10 minutes to ~2.5 hours. During head fixation acclimatisation sessions mice were also given 15% sucrose solution. The drinking interval was ~15 min in duration, with a spout providing a slow continuous flow of liquid (33.3 μl/min × 15 min = 0.5 ml) to the mouse. The consumed volume was established by collecting and measuring the volume of liquid that was not consumed and subtracting it from the total extruded volume of 0.5 ml. Once the animals were acclimatised to head-fixation and drinking the sucrose solution, the solution was adulterated with either quinine hydrochloride (0.5 mM) or sucrose octoacetate (0.5 mM) bitterants.

Fully acclimatised mice were able to drink up to ~0.5 ml of the bitter-sweet liquid (1 mg/ml TCB-2, 0.5 mM bitterant, 15% sucrose, w/v). We proceeded to record neural activity from a mouse once it reliably drank at least 0.15 ml during training and had reached the full 2.5 h duration of head-fixation acclimatisation.

### Craniectomies

Craniectomies were performed once animals were fully acclimatised to head-fixation and drinking of bitter-sweet liquid. Craniectomies were performed in aseptic conditions under general isoflurane anaesthesia, with the animal head-fixed using the previously implanted headplate. Small hole was drilled in the skull at previously marked coordinates above medial prefrontal cortex (+1.8mm anterior, 0.5mm lateral to Bregma). Craniectomies were protected with Kwik-Cast before and between recordings. For recovery from anaesthesia, the animal cage was placed on top of a heated recovery shelf for at least 1 hour.

### Electrophysiological recordings and spike-sorting

Neuropixels probes were lowered through the dura to a depth of 3-3.5 mm at 2 μm/s and allowed to settle for 10 minutes prior to the beginning of the recording. Signals from the Neuropixels 1.0 probes were acquired using Spike GLX software (http://billkarsh.github.io/SpikeGLX) and saved into a high-pass filtered (0.3-10 kHz) signal from each recording site at 30 kHz resolution and a low-pass filtered (0.5-1000 Hz) LFP signal at 2.5 kHz. The latter signal was used for LFP analysis (Figure 1D; 2 recordings with excessive noise in the LFP frequency band were excluded from the analysis). The former signal was used for spike-sorting, which was performed in two stages. First, automatic spike sorting was performed using KiloSort (Pachitariu et al., 2016) (versions 1 or 2). The results were manually curated using Phy (https://github.com/cortex-lab/phy). We obtained 256±138 well isolated units per recording (mean and st.dev.).

To verify that our main findings concerning the differences between TCB-2 and control recordings (figures 2–4) do not depend on the manual curation, the recordings were also sorted using KiloSort 2.5 and the analyses were repeated on all units that were automatically tagged as ‘good’ (while discarding all ‘mua’ units), thus eliminating the manual curation step. The quantitative results were similar to the manually curated data.

### Pupil imaging

To image the pupil during the electrophysiological recordings we used a USB camera (Full HD Webcam, ELP, China), with a 50 mm focal length M12 mount lens (ACM12B5025IRM12MM, AICO Electronics Limited, China), which was attached to the camera using several spacer rings. The camera was positioned ~15 cm from the head of the animal, providing a field of view of ~8×8 mm. A custom script in MATLAB saved the pupil video (at 24 frames/s) and generated a signal for synchronisation with the electrophysiological recording.

Pupil videos were processed in two batches using the open-source DeepLabCut software (Mathis et al., 2018). 20-40 frames were extracted from each video to form two batches of 300 and 175 frames. Using the annotation GUI, in each frame we marked the pupil edge at four points, 90° apart. For each batch, the deep neural networks were trained with 90% of the labelled frames and tested with 10%. The trained networks were used to determine the position of the four pupil edge markers for each frame of each video. Opposing markers were taken to define the two axes of an ellipse, whose area was taken as the pupil size.

### Histology

In a subset of recordings the Neuropixels 1.0 probe was coated with DiI (Invitrogen) to visualise the probe tract (as shown in Figure 1B). At the end of the recording, animals were deeply anaesthetised with pentobarbital (50 mg/kg, i.p.) and transcardially perfused with phosphate buffered saline and then paraformaldehyde (5 minutes each). The brain was removed and stored at 4° C in 4% paraformaldehyde for 24 hours, with 30% sucrose (w/v) added subsequently. Coronal 80 μm slices were sectioned on a freezing microtome (Rankin Biomedical Corporation), then mounted and stained with DAPI (Vectashield). Slices were imaged using a DM2500, Leica microscope.

### Data analysis

Data was analysed using custom functions and scripts implemented in MATLAB R2020a.

For the step function fit (Figure 2), the firing rate of each neuron in the interval [-19.5, +48.5] minutes around the time point at which the spout was made available to the mouse was represented as a signal at 1 Hz resolution. This signal was approximated using a step function that minimises the total residual error, which involves finding a point at which splitting the signal into two parts minimises the sum of the residual squared error of each part from its own mean. This was performed using MATLAB’s `findchangepts` function.

Analysis of firing rate distributions across an entire population (or subpopulation) proceeded as follows. First, we excluded units with a firing rate < 0.01 spikes/s. Next, the firing rates were fitted by a gamma distribution (using MATLAB’s `gamfit` function). We used gamma rather than log-normal distribution because we consistently found that it provided fits with better log-likelihood (Figure S1). Once the two parameters specifying the best gamma fit (*a*, *b*) were found, the bell-shaped analytical distribution of the log-rates *r* (as plotted in Figures 4–6) is specified by the probability density function of the log-gamma distribution, which (for base 10 logarithm) is given by the following equation:

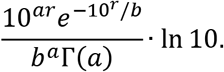

Log-rates of a neuronal population in two different brain states were compared by their means and variances. The means were compared by a two-sample t-test that does not assume equal variance (which relied on the fact that the log-rates, while somewhat skewed, were approximately normally distributed). The variances were compared by Levene’s quadratic test.

Firing rate of a neuron within a particular brain state cannot be measured with absolute precision from any finite sized data, and particularly so when the available data is only several minutes in duration. Denoting the true log-rate of neuron *i* in brain states *Y* and *Z* by 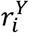 and 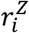, our approximate estimate will be 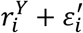 and 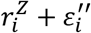, where 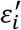 and 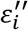 are the measurement error factors. If the very same data is used to classify the neurons into the two subpopulations and then to test for difference within the subpopulation across the brain states, entirely spurious results can be obtained. As a simple example, consider the case of firing rates that are independent of brain state (i.e., 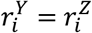 for every *i*). Then, neurons for which 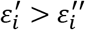 are assigned to the “*Y* → *Z* downregulated” subpopulation, and neurons for which 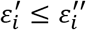 are assigned to the “*Y* → *Z* upregulated” subpopulation. Statistical tests comparing the distribution of 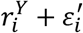 to the distribution of 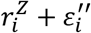 within each subpopulation might show highly significant differences in the mean or variance, which however are entirely spurious (and such a scenario can be easily numerically simulated). To avoid this problem, we performed cross-validation by using non-overlapping parts of the data for the assignment of neurons to subpopulations and for the comparison across brain states. Specifically, once the assignment to the two subpopulations was done as described above, the statistical tests were performed on the distributions of 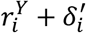 and 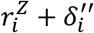, where 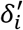 and 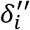 are the error factors in measuring the true log-rates from a different and non-overlapping part of the recording (and thus 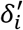 and 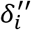 are presumed to be independent from 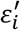 and 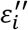). In particular, if firing rates are independent of brain state, no spurious significant differences should be found.

### Allen Brain Institute data

We have analysed data of Neuropixels recordings in mice publicly released by the Allen Brain Institute (Siegle et al., 2021). We relied for the analyses on 30 minutes of spontaneous activity (when nothing was shown on the screen in front of the animal) recorded from each animal, along with their pupil size and running speed, as part of a much longer recording session. For the analysis we used only intervals where the mouse was not locomoting, and we discarded recordings where less than 3 minutes (of the 30 minutes) were in this condition. Thus, we have used data from 20 animals (not all brain areas were recorded in each animal). Here, we limited our analysis to neurons in the primary visual cortex (VISp), area CA1 of the hippocampus and the dorsal lateral geniculate nucleus (LGd).

## Supporting information

Supplemental Video (Head twitch response)

## Author contributions

M.O. designed the study; B.D. performed the experiments with assistance from M.D. and M.O.; B.D. and M.O. wrote the paper; all authors contributed to data analysis and editing.

## Acknowledgements

We thank James McCutcheon for advising us on the available bitterants during the early stages of the project, and Todor Gerdjikov and John Apergis-Schoute for their comments on an earlier draft. This study was supported by the Academy of Medical Sciences and Wellcome Trust (Springboard award SBF002\1045) and BBSRC (grant BB/P020607/1 and PhD studentship 2265967).

## Supplementary materials

**Figure S1.**
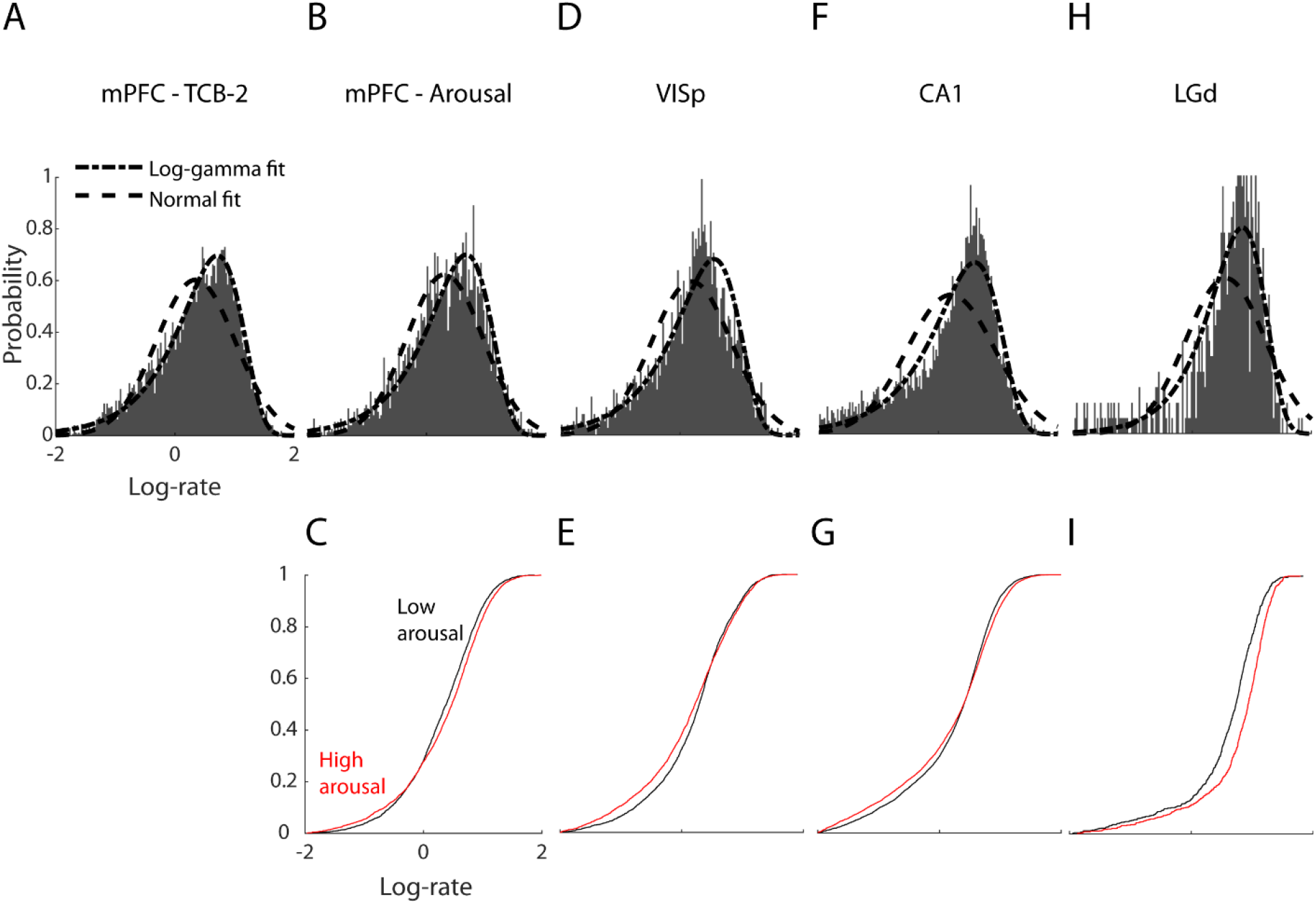
Distribution of log firing rates and analytical fitting. **(A)** Probability distribution of log firing rates in mPFC pooled across control and TCB-2 recordings before drinking (7176 neurons in total), with log-gamma fit (log-likelihood/sample = −2.60) and normal fit (log-likelihood/sample = −2.64). The log-gamma analytical distribution provided a better fit to the empirical data. **(B)** Probability distribution of log firing rates in mPFC in low arousal condition, with log-gamma fit (log-likelihood/sample = −2.50) and normal fit (log-likelihood/sample = −2.51), same scale as (A). **(C)** Cumulative distribution plots of mPFC log-rates in low- and high-arousal states. The high arousal distribution (shown in red) is visibly wider (its left tail is shifted to the left and the right tail is shifted to the right). **(D, E)** Same format as (B, C) for log firing rates in VISp, log-gamma fit had log-likelihood/sample = −2.26 and normal fit had log-likelihood/sample = −2.29. **(F, G)** Same format as (B, C) for log firing rates in CA1, log-gamma fit had log-likelihood/sample = −2.35 and normal fit had log-likelihood/sample = −2.45. **(H, I)** Same format as (B, C) for log firing rates in LGd, log-gamma fit had log-likelihood/sample = −2.93 and normal fit had log-likelihood/sample = −3.13.

**Figure S2.**
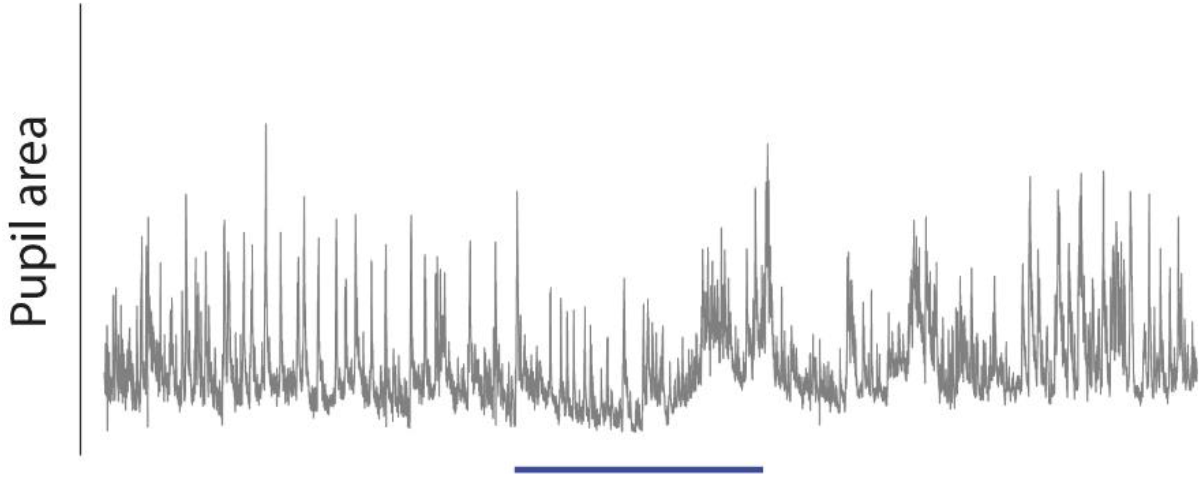
Pupil area in a representative recording. The blue bar indicates a 15-minute interval during which liquid with TCB-2 was available through a drinking spout.

**Figure S3.**
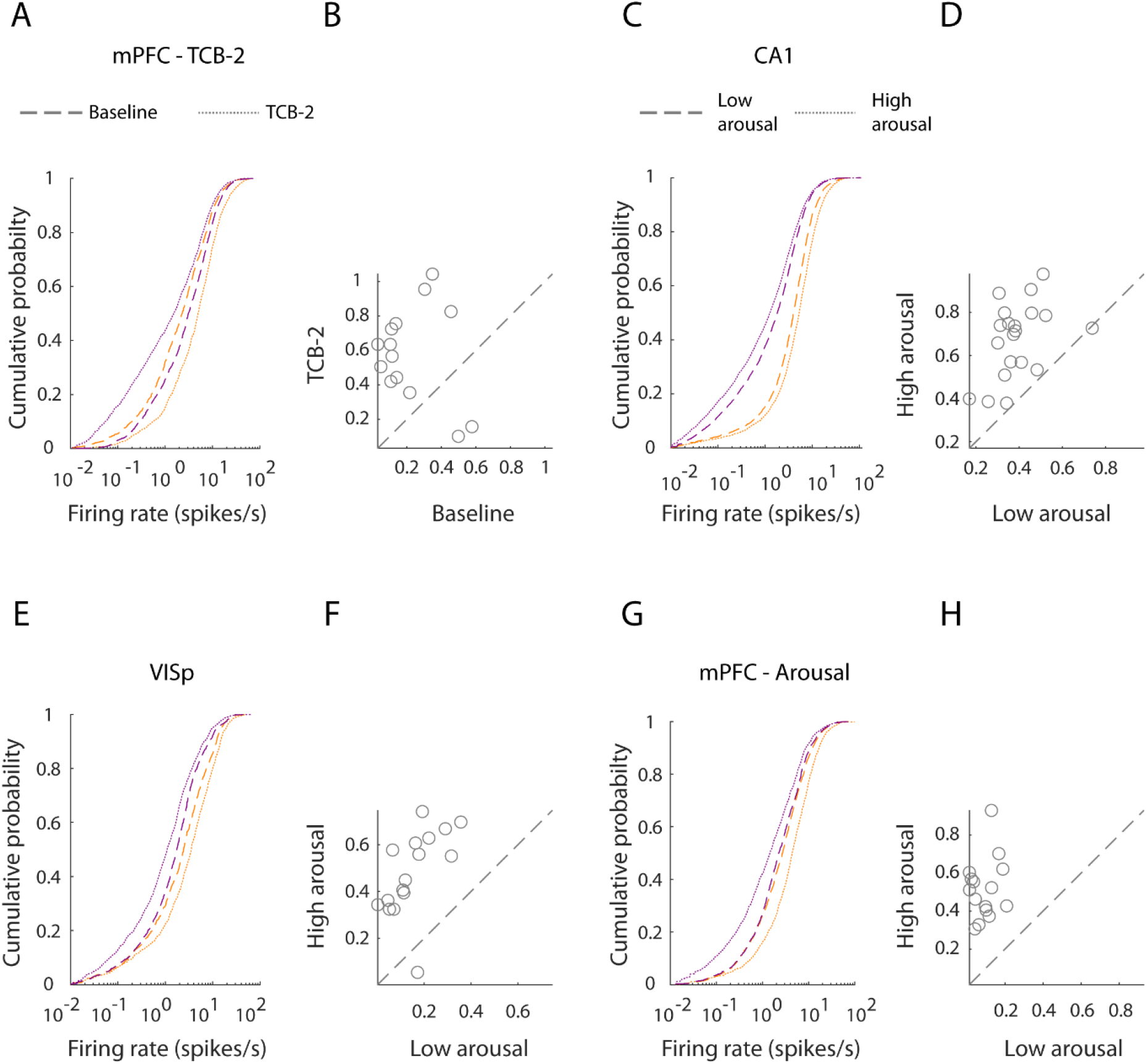
Changes in the firing rates of the upregulated and downregulated subpopulations in psychedelic and arousal-driven brain state transitions. **(A)** Cumulative distribution of firing rates of subpopulations of mPFC neurons that elevated (orange) and supressed (purple) their firing rate as a result of TCB-2 administration (3564 neurons, 14 recordings). **(B)** Distance between log firing rates means of the two subpopulations before and after ingesting TCB-2 (P = 2.8×10^−3^, t-test). **(C, D)** Same format as (A, B) for CA1 subpopulations upregulated and downregulated by arousal changes in low (dashed line) and high (dotted line) arousal states (P = 2.2×10^−7^, 7729 neurons, 20 recordings). **(E, F)** Same format as (C, D) for VISp data (P = 7.5×10^−8^, 2831 neurons, 18 recordings). **(G, H)** Same format as (C, D) for mPFC data (P = 1.1×10^−7^, 3100 neurons, 15 recordings).

**Figure S4.**
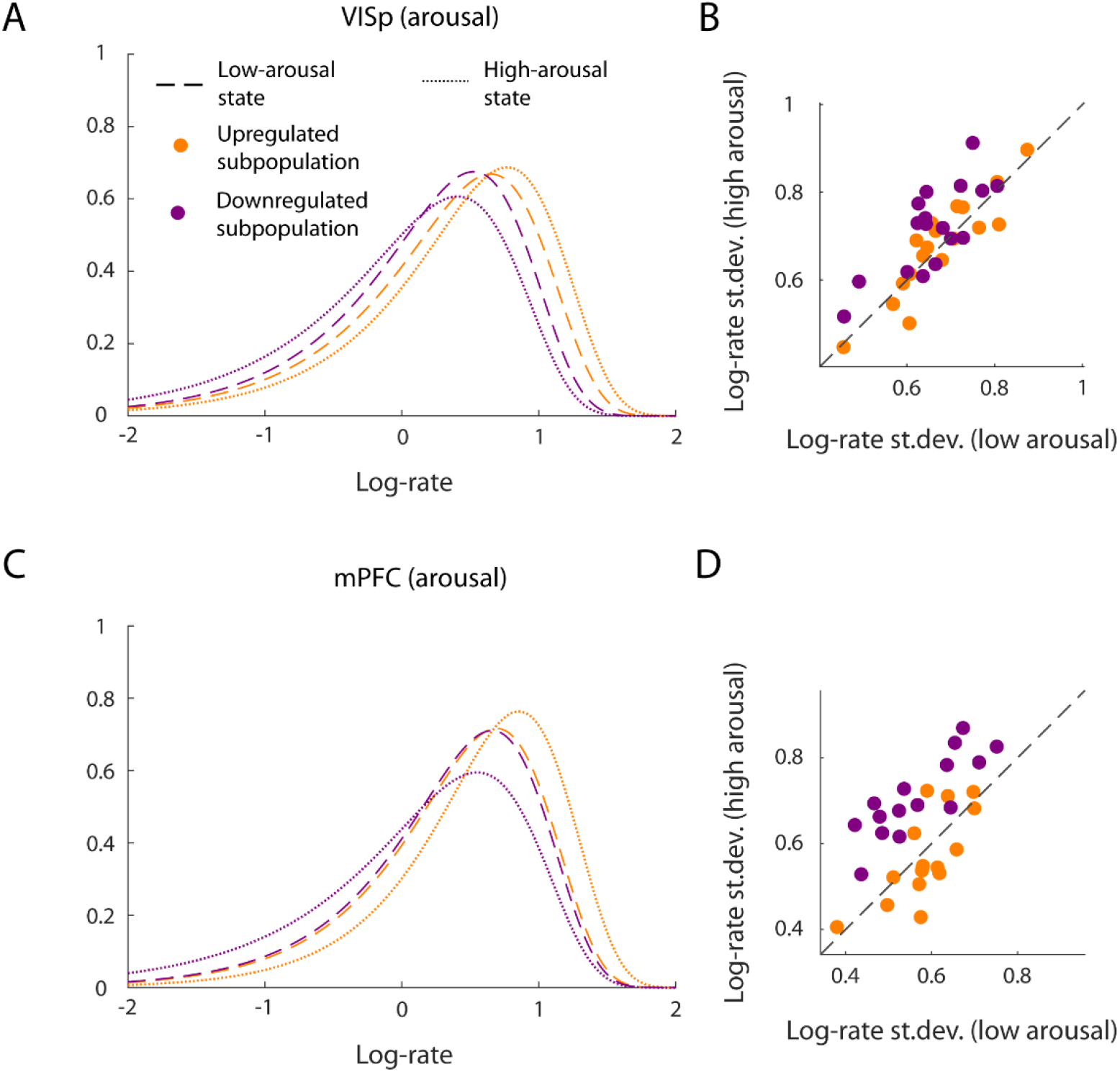
Stretching and squeezing of the log-rate distributions of the upregulated and downregulated subpopulations. **(A)** Log-gamma fit of the distribution of log-rates of VISp neurons whose firing rates were upregulated by increased arousal (41%, orange) and of neurons whose firing rates were downregulated by increased arousal (59%, purple) in low- (dashed line) and high-arousal (dotted line) brain states. **(B)** Standard deviation of the VISp log-rates in low- and high-arousal brain states of subpopulation upregulated (orange) and downregulated (purple) by arousal. In the former subpopulation, changes in arousal did not change the st.dev. of the log-rate distribution at the level of individual recordings (low: 0.68±0.10 vs high: 0.70±0.13, P = 0.23, t-test of the st.dev. in low and high arousal, 18 recordings). In the latter subpopulation, high arousal led to an increase of st.dev. of the log-rate distribution (low: 0.66±0.09 vs high: 0.72±0.10, P = 1.8×10^−3^, t-test of the st.dev. in low and high arousal). **(C, D)** Same format as (A, B) for mPFC. In the subpopulation upregulated by increase in arousal, changes in arousal did not change the st.dev. of the log-rate distribution (low: 0.59±0.07 vs high: 0.58±0.10, P = 0.60, t-test of the st.dev. in low and high arousal, 15 recordings). In the subpopulation downregulated by increase in arousal, high arousal led to an increase of st.dev. of the log-rate distribution (low: 0.57±0.10 vs high: 0.71±0.09, P = 1.6×10^−7^, t-test of the st.dev. in low and high arousal).

**Supplementary Video.**
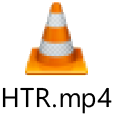
Head Twitch Response. An example of a single head twitch bout observed after the animal was given TCB-2. The video is slowed down to ×0.66. The HTR is in the second half of the clip.

